# Identification of Common Molecular Signatures Shared between Alzheimer’s and Parkinson’s Diseases and Therapeutic Agents Exploration: An Integrated Genomics Approach

**DOI:** 10.1101/2020.12.31.424962

**Authors:** Nairita Ahsan Faruqui, Durdana Hossain Prium, Sadrina Afrin Mowna, Tanjim Ishraq Rahaman, Arundhati Roy Dutta, Mst. Farjana Akter

## Abstract

Alzheimer’s disease (AD) and Parkinson’s disease (PD) are two most prevalent age-related dementias that severely affect a large number of elderly people around the globe. Poor understanding of pathogenesis of these neurological diseases imposes challenge to discover therapeutic measures and effective diagnosis methods. In this study, a network-based approach was utilized to identify potential common molecular signatures and therapeutic agents for AD and PD. Protein-protein interaction analysis revealed NCK1, UBC, CDH1, CDC20, ACTB, PSMA7, PRPF8, RPL7, XRCC6 and HSP90AB1 as the best proteome signatures. Different regulatory transcriptional signatures i.e., YY1, NFKB1, BRCA1, TP53, GATA2, SREBF2, E2F1, FOXC1, RELA and NFIC and post-transcriptional signatures i.e., hsa-mir-186-5p, hsamir-92a-3p, hsa-mir-615-3p, hsa-let-7c-5p, hsa-mir-100-5p, hsa-mir-93-3p, hsa-mir-5681a, hsamir-484, hsa-mir-193b-3p and hsa-mir-16p-5p were identified from other interaction network. Drug-gene interaction study revealed possible therapeutic agents which may reverse the AD and PD condition. The scientific approach of this study should contribute to identify potential biomarkers, drug targets and therapeutic agents against AD and PD which should in turn advance the present efforts of scientists to secure effective diagnosis and therapeutic options. However, further *in vivo* and *in vitro* experiments might be required to validate the outcomes of this study.

## Introduction

Neurodegenerative disease is an umbrella term for a range of conditions which primarily affect the nerve cells in the brain. Among the several neurodegenerative diseases, Alzheimer’s disease (AD) and Parkinson’s disease (PD) are the most prevalent among the mass population of the world [1]. About 44 million people are diagnosed with AD worldwide, and about 10 million are affected with PD. AD is characterized by the accumulation of protein aggregates of amyloid and neurofibrillary tangles containing hyperphosphorylated neuronal tau protein. It is a form of dementia that includes slow deterioration of thinking, behavioral skills and loss of memory, most likely to affect people over 65 years of age [2][3]. Genetics, head injuries, depression and hypertension are possible risk factors which may also contribute to an early onset of the disease. The symptoms of this progressive disorder worsen over time and leads to difficulties with language, disorientation, mood swings, lack of motivation and self-care.

Even though the disease is affecting a large number of people around the world, there is currently no effective diagnosis or therapeutic strategies available. Due to the lack of any definitive diagnosis method of Alzheimer’s disease, apart from physical assessments and a review of the medical history, cognitive testing, neurological exams, blood tests or brain imaging scans (CT, MRI, PET, etc.) can be used as probable assessment methods [4]. Brain imaging methods enable doctors to only detect specific brain changes caused by the disease, but the information obtained from these tests does not provide any significant value to making a proper diagnosis for the disease. Examination of brain tissues in autopsies can be used to accurately detect Alzheimer’s only after the death of the individual.

There are no effective treatments available for Alzheimer’s, although there are few possible drugs available for AD, they do not necessarily cure the disease or stop its progression, and are also known to wear off over time. The drug, Cholinesterase inhibitor (mild to moderate AD) and Memantine (moderate to severe AD) can be used to treat the symptoms but cannot reverse Alzheimer’s or cure the disease and thus, are known to be less effective [5][6]. On the other hand, these drugs have severe side effects and the effect of Cholinesterase inhibitors eventually wears off, along with limitations which may include its inability to be used by patients of cardiac arrhythmias. Moreover, the established treatments for AD are mainly symptomatic in nature which focuses mainly on counterbalancing the neurotransmitter disturbance of the disease [7]. Again, treatment for AD requires drugs to bypass the blood brain barrier (BBB) which allows only selective molecules to enter the brain [8]. Therefore, the restriction of the barrier to transport the drug to the brain limits the availability of drugs for Alzheimer’s.

PD, on the other hand, is a progressive degenerative disorder of the central nervous system that mainly affects the motor system of our brain [9]. The symptoms of Parkinson’s are known to emerge slowly, and non-motor symptoms are seen as the disease progresses and worsens. The symptoms include tremors, trouble with thinking and behavior, depression, anxiety, sensory, sleep and other emotional problems. Cell death in the substantia nigra, a region in the midbrain, is responsible for the motor symptoms of the disease which also involves the accumulation of proteins into Lewy bodies in the neurons [10]. Parkinson’s disease typically occurs in individuals over the age of 60. Diagnostic tests such as neuroimaging are used in the diagnosis of typical cases, mainly based on symptoms, along with the ruling out of other diseases [11]. Although there is no cure for Parkinson’s, the treatments are aimed at improving symptoms. The treatment is usually initiated with the antiparkinsonian medication levodopa (L-DOPA), followed by dopamine antagonists when L-DOPA becomes less effective. With the progression of the diseases and neuronal loss, the medications show a lack of effectiveness alongside contributing to complications marked by writhing movements. The progression of the symptoms emerges due to the course of cell loss in Parkinson’s and at the time of detection, almost around half of the dopamine producing brain cells are seen to be lost from the substantia nigra [12]. The existing Parkinson drugs only aim to replace or mimic the action of dopamine in the brain in order to mask the progressing symptoms. The continued loss of brain cells requires an increased use of these drugs which contribute to the following side effects of these medications, that impact the quality of life of the individual drastically. Since the existing diagnosis methods do not provide specific and credible evidence for the detection of either Alzheimer’s or Parkinson’s, biomarkers can be used as probable indicators for early diagnosis of these diseases [13]. The early diagnosis of Alzheimer’s and Parkinson’s will help keep the effects of the diseases on the patient at bay. In light of this, biomarkers are useful tools that may contribute to the early detection of these diseases [14]. And the identification of new molecular targets and therapies to treat AD and PD continues to be the major race of time for the scientists.

In our study, we sought to explore the common molecular signatures, drug targets and therapeutic candidate molecules for AD and PD that include proteins, transcription factors and miRNAs utilizing publicly available microarray data and applying network-based integrated approach (**Figure 1**). Then the identified signatures were further analyzed to check for any contribution in the pathogenesis of AD, PD and other diseases.

**Figure 1:**
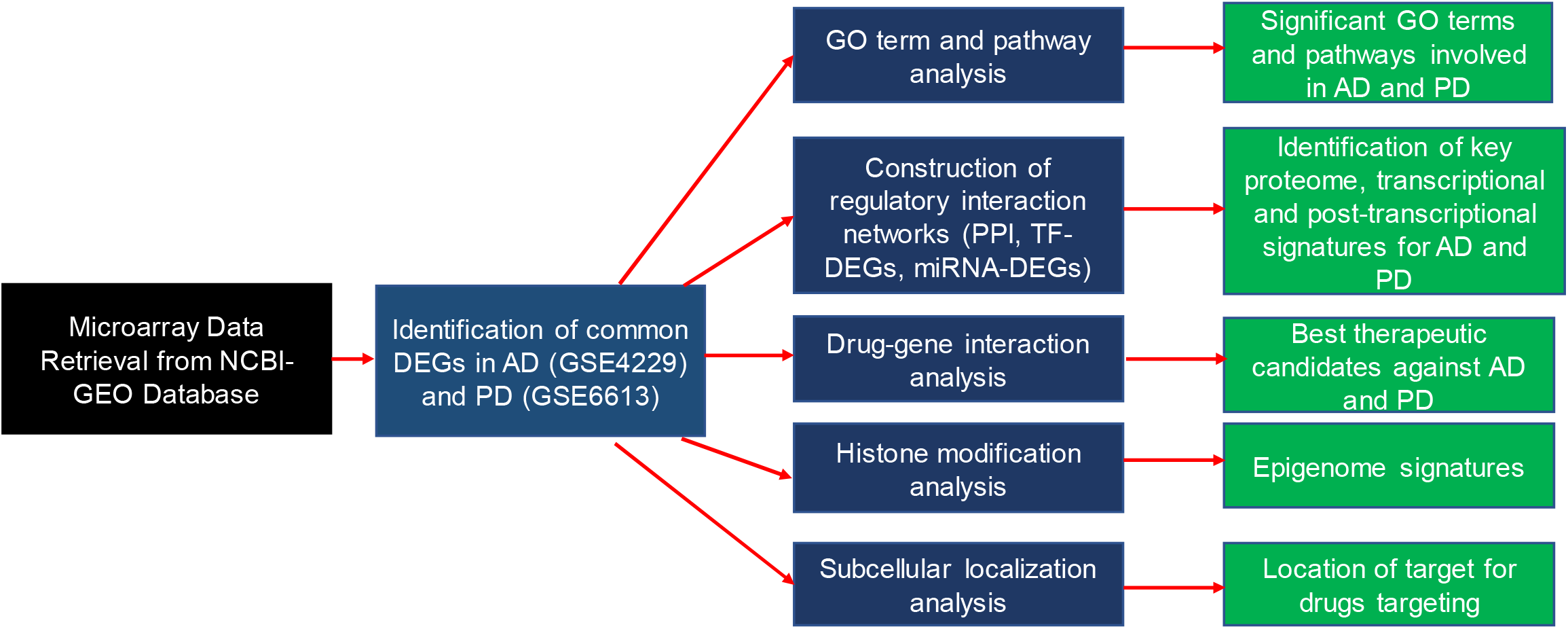
Strategies employed in overall study. Two different microarray datasets for AD (GSE4229) and (GSE6613) were retrieved and common DEGs in AD and PD were identified. Then the DEGs were analyzed to identify potential proteome, transcriptional and post-transcriptional signatures. The DEGs were also analyzed for their pattern of histone modification and subcellular localization. The drug-gene interaction was carried out to find potential therapeutic agents to be triggered against AD and PD. AD: Alzheimer’s disease; PD: Parkinson’s disease; DEGs: differentially expressed genes; TF: Transcription factor; PPI: Protein-protein interaction network.

Genomic, proteomic and other omic-based approaches are now being widely used in different biomedical researches for the optimization of disease mechanism, identification of molecular targets and biomarkers for specific diseases [15]. The application of network-based or pathway-based strategy to omic data helps to understand biological processes, gene regulatory networks, functional relations of genes and proteins which then drives the discovery of biomarkers, molecular targets and therapeutics for a particular disease [16] [17]. The scientific findings of this study will help to discover effective biomarkers and molecular targets for drugs to be triggered against AD and PD.

## 2. Materials and Methods

### 2.1. Data Retrieval and Differential Gene Expression Analysis

The microarray data of AD and PD patient was retrieved from Gene Expression Omnibus (GEO) database of National Centre for Biotechnology Information (NCBI); NCBI-GEO. GSE4229 microarray data containing total RNA profile of peripheral blood of 22 normal elderly control and 18 AD patients, was collected for analysis [18]. GSE6613 was selected as the sample data for PD patient which contains whole blood expression data of 50 PD patients, 23 healthy normal control and 33 neurodegenerative patients other than PD [19]. Only the PD patient (n=50) and healthy normal control (n=23) samples were analyzed for differential gene expression pattern analysis in PD patient. Differentially expressed genes (DEGs) were identified using the GEO2R tool of NCBI, where at first, Log2 transformation was applied and then Benjamini & Hochberg correction method was applied to control the false discovery rate. The < 0.05 P value cut off criterion was used to identify the significant DEGs for both datasets. Then the overlapping DEGs of 2 samples were selected using the online tool InteractiVenn [20].

### 2.2. Functional Enrichment Analysis of DEGs

Gene co-expression, pathway and interaction network analysis allows the identification of functionally correlated genes and disease-gene interaction and understanding of their tentative function and regulatory biomolecules [21] - [23]. Afterward, the identified overlapping DEGs were then subjected to analyze their gene ontology (GO) terms in order to understand their biological processes, molecular functions and as part of different cellular components using the Enrichr. PANTHER online tool was used to generate the protein class over-representation [24] [25]. The < 0.05 P value cut off was considered as the selection criterion of the ontology terms. Different pathways involved with the selected DEGs were also analyzed with the similar platform.

### 2.3. Protein-Protein Interaction (PPI) Network and Hub Protein Analysis

Analysis of the transcriptome provides significant information about the proteome of an organism leading to the prompt identification of molecular targets and signatures. PPI interaction networks holds substantial information about the key proteins which can drive the discovery of biomarker signatures and drug targets [26]-[28]. Hub proteins represent the most connected nodes in a PPI network and they provide useful insight about the function and key proteins of any network [29]. The generic PPI network was constructed using the STRING database using a high confidence score of 900 [30]. Thereafter, the PPI network was analyzed in Cytoscape and Hub proteins (top 10 mostly connected nodes) were identified with the cytoHubba plugin utilizing the betweenness centrality matrix [31] [32]. Afterward, the identified Hub proteins were analyzed for their involvement in different pathways using Panther database.

### 2.4. Identification of Regulatory Biomolecules

Transcription factors (TFs) and microRNAs (miRNAs) regulate the expression of many genes and thus these two biomolecules provide two significant sources of molecular targets [33]. Many non-coding RNA and TFs-based biomarkers have been approved for different diseases including diabetes and cancer [34] [35]. Hub gene were analyzed against JASPAR which is an open access, curated and non-redundant database of DNA binding transcription factors, with the help of NetworkAnalyst to construct transcription factor (TF)-Hub genes interaction network [36] [37]. miRNA-Hub gene interaction network was created searching the hub proteins against TarBase database, a manually curated microRNA database which includes almost 1,300 experimentally supported targets [38]. Top 10 interacting nodes (TFs and miRNAs) with most edges were selected and analyzed.

### 2.5. Identification of Epigenome Signatures

The pattern of histone modification including acetylation and methylation influences the expression of specific genes. These modifications provide profound information about the onset and progression of particular disease [39] [40]. The identified hub genes were analyzed for their potential histone modification sites using the Epigenomics Roadmap Chip-seq dataset with the help of Enrichr [41]. P value <0.05 filter was used as the selection criterion.

### 2.6. Cross Validation of the Identified Signatures

The selected hub proteins and TFs were searched against MalaCards human disease database. MalaCards is an integrated and comprehensive database which collects information on dysregulated genes, proposed signatures and diseases from more than 70 sources including updating literatures [42]. All of the proteome signatures were found to be involved in multiple genetic diseases and disorders. Different proteomic signatures were found to be involved and proposed as signatures of different neurological diseases and mental complications i.e., AD, Huntington’s disease, bipolar disorders, dementia, neuroretinitis, mental depression, autism spectrum disorder, glioma, glioblastoma, schizophrenia etc. Different studies predicted the dysregulation of different identified miRNAs involved in the prognosis of neurological diseases i.e, AD, PD, HD, bipolar disorders [43]-[46].

### 2.7. Identification of Therapeutic Candidate Molecule

The identified hub genes and selected transcription factor encoding genes were searched against the drug-gene interaction database (DGIdb) to identify potential therapeutic candidate molecules. DGIdb contains information of almost 10,000 drugs involved in 15,000 drug-gene interaction or belonging to one of total 39 potentially druggable gene categories [47]. Different selected hub genes and TF encoding genes were found to interact with different drug molecules. Best candidate molecules for a gene were selected based on the drug scores.

### 2.8. Determination of Subcellular Localization

Since the role of a particular protein depends on the subcellular localization of that protein, it is vital to determine the localization in order to understand the protein function [48]. The selected hub proteins were analyzed with the online tool-WoLF PSORT to determine their subcellular localization [49].

## 3. Results

### 3.1. Transcriptome Signatures in AD and PD

After retrieval the publicly available microarray was statistically analyzed to identify potential DEGs in AD and PD patients. The DEGs were then subjected to further experiment to select common DEGs in AD and PD which were then utilized in this experiment. A total of 183 common DEGs were selected from the two different datasets (**Figure 2A**). Then the classes of the expressed proteins by the DEGs were analyzed. The DEGs were found to be involved in predominantly expressing translational proteins (24%), metabolite interconversion enzymes (16%), protein modifying enzymes (14%), and nucleic acid binding proteins (13%) (**Figure 2B**).

**Figure 2:**
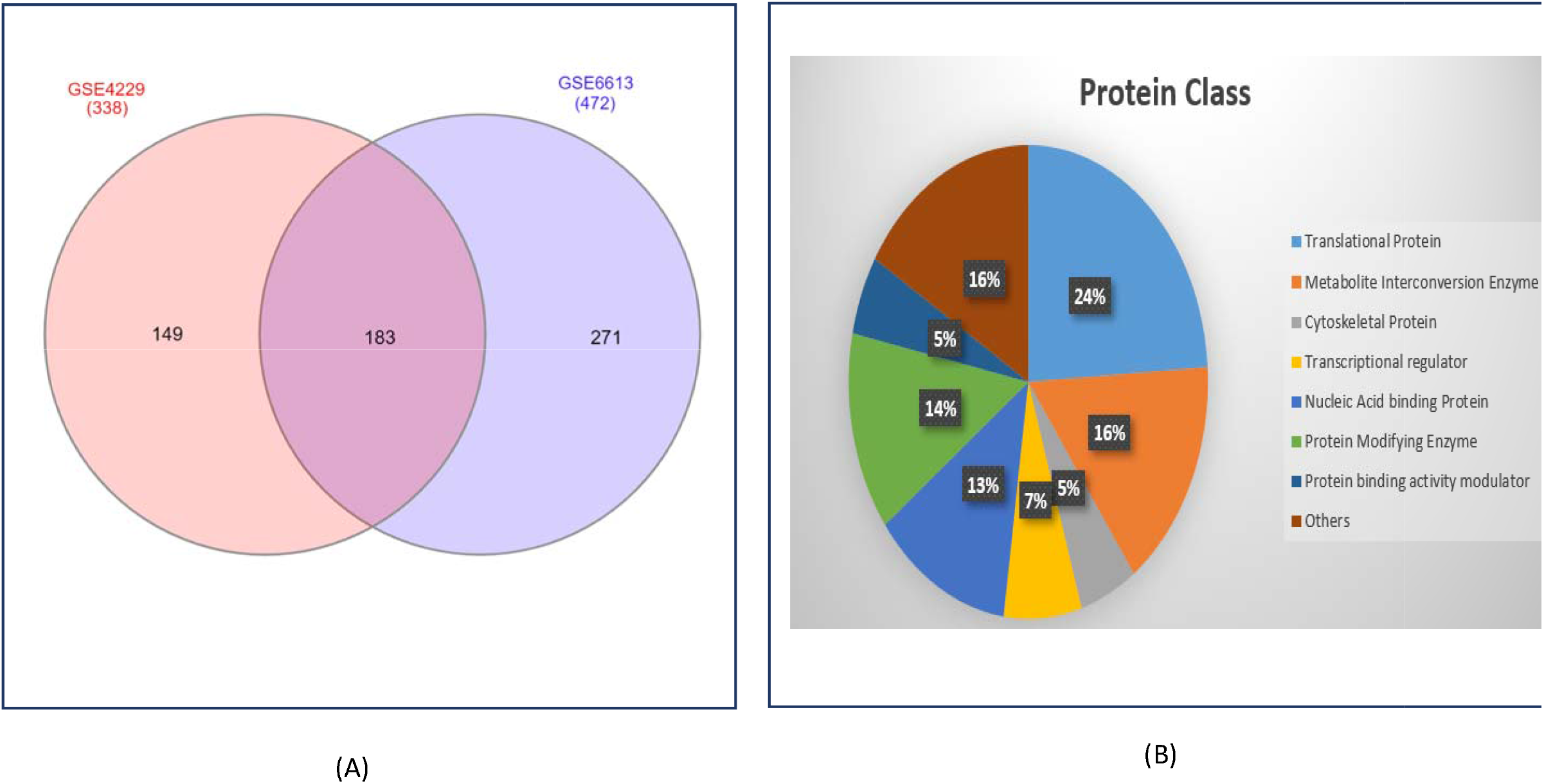
(A) Ven diagram of microarray datasets of AD (GSE4229) and PD (GSE6613) patients. (B) Protein class overrepresentation of the identified common DEGs in AD and PD.

Then the DEGs were analyzed for their involvement in pathways and evidence of their involvement in different pathways was recorded (**Table 1**). Enrichment analysis of the DEGs showed that the genes were involved in different biological processes, molecular function and cellular components. This analysis revealed that the common DEGs were primarily involved in ribosome mediated function and protein processing and trafficking (**Table 2**).

**Table 1:** Different pathways involved with different DEGs in AD and PD.

| Pathway | Type | Adjusted P Value | Genes Involved |
| --- | --- | --- | --- |
| BioPlanet | Cytoplasmic ribosomal proteins | 2.993e-8 | RPL5, RPL31, RPL0, RPL23, RPS20, RPL24, RPL37A, RPL36A, RPS25, RPS13 |
| KEGG | Ribosome | 3.367e-9 | RPL5, RPL31, RPL0, RPL23, RPS13, RPL24, RPL36AL, RPL6, MRPS5, RPS3A, MRPL27 |
| BioCarta | Downregulated of MTA-3 in ER-negative Breast Tumors Homo sapiens h mta3 Pathway | 0.01815 | CDH1, ALDOA, GAPDH |
| Reactome | Viral mRNA Translation Homo sapiens | 2.334e-8 | RPL5, RPL31, RPL0, RPL23, RPS13, RPL24, RPL6, RPS3A, RPS25, RPLP0, RPS13, RPL36A, RPL37A |

**Table 2:**
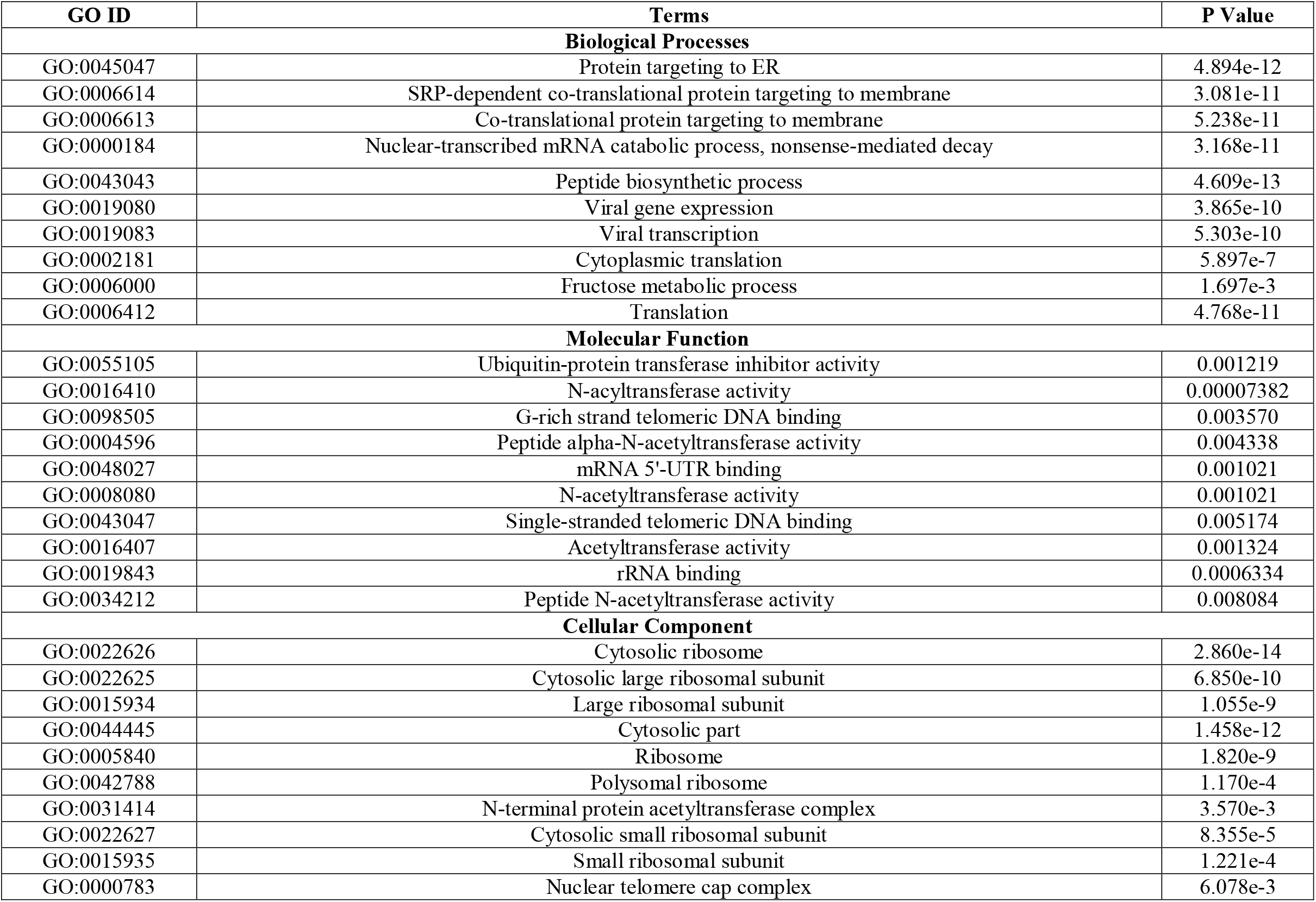
Best ten selected gene ontology (GO) terms of the selected common DEGs in AD and PD.

| GO ID | Terms | P Value |
| --- | --- | --- |
| **Biological Processes** |  |  |
| GO:0045047 | Protein targeting to ER | 4.894e-12 |
| GO:0006614 | SRP-dependent co-translational protein targeting to membrane | 3.081e-11 |
| GO:0006613 | Co-translational protein targeting to membrane | 5.238e-11 |
| GO:0000184 | Nuclear-transcribed mRNA catabolic process, nonsense-mediated decay | 3.168e-11 |
| GO:0043043 | Peptide biosynthetic process | 4.609e-13 |
| GO:0019080 | Viral gene expression | 3.865e-10 |
| GO:0019083 | Viral transcription | 5.303e-10 |
| GO:0002181 | Cytoplasmic translation | 5.897e-7 |
| GO:0006000 | Fructose metabolic process | 1.697e-3 |
| GO:0006412 | Translation | 4.768e-11 |
| **Molecular Function** |  |  |
| GO:0055105 | Ubiquitin-protein transferase inhibitor activity | 0.001219 |
| GO:0016410 | N-acyltransferase activity | 0.00007382 |
| GO:0098505 | G-rich strand telomeric DNA binding | 0.003570 |
| GO:0004596 | Peptide alpha-N-acetyltransferase activity | 0.004338 |
| GO:0048027 | mRNA 5'-UTR binding | 0.001021 |
| GO:0008080 | N-acetyltransferase activity | 0.001021 |
| GO:0043047 | Single-stranded telomeric DNA binding | 0.005174 |
| GO:0016407 | Acetyltransferase activity | 0.001324 |
| GO:0019843 | rRNA binding | 0.0006334 |
| GO:0034212 | Peptide N-acetyltransferase activity | 0.008084 |
| **Cellular Component** |  |  |
| GO:0022626 | Cytosolic ribosome | 2.860e-14 |
| GO:0022625 | Cytosolic large ribosomal subunit | 6.850e-10 |
| GO:0015934 | Large ribosomal subunit | 1.055e-9 |
| GO:0044445 | Cytosolic part | 1.458e-12 |
| GO:0005840 | Ribosome | 1.820e-9 |
| GO:0042788 | Polysomal ribosome | 1.170e-4 |
| GO:0031414 | N-terminal protein acetyltransferase complex | 3.570e-3 |
| GO:0022627 | Cytosolic small ribosomal subunit | 8.355e-5 |
| GO:0015935 | Small ribosomal subunit | 1.221e-4 |
| GO:0000783 | Nuclear telomere cap complex | 6.078e-3 |

### 3.2. Proteome Signatures in AD and PD

The common DEGs were then utilized to construct a generic PPI network from which 10 most connected key proteins (Hub proteins) were selected. NCK1, UBC, CDH1, CDC20, ACTB, PSMA7, PRPF8, RPL7, XRCC6 and HSP90AB1 were identified as the most connected nodes or hub proteins from the PPI network which were then analyzed in different phases of this experiment (**Figure 3 and Table 3**). The selected hub proteins were then also analyzed for their involvement in key pathways. Significant involvement of few proteins with Alzheimer’s disease-presenilin pathway and Parkinson’s disease was recorded (**Table 4**).

**Figure 3:**
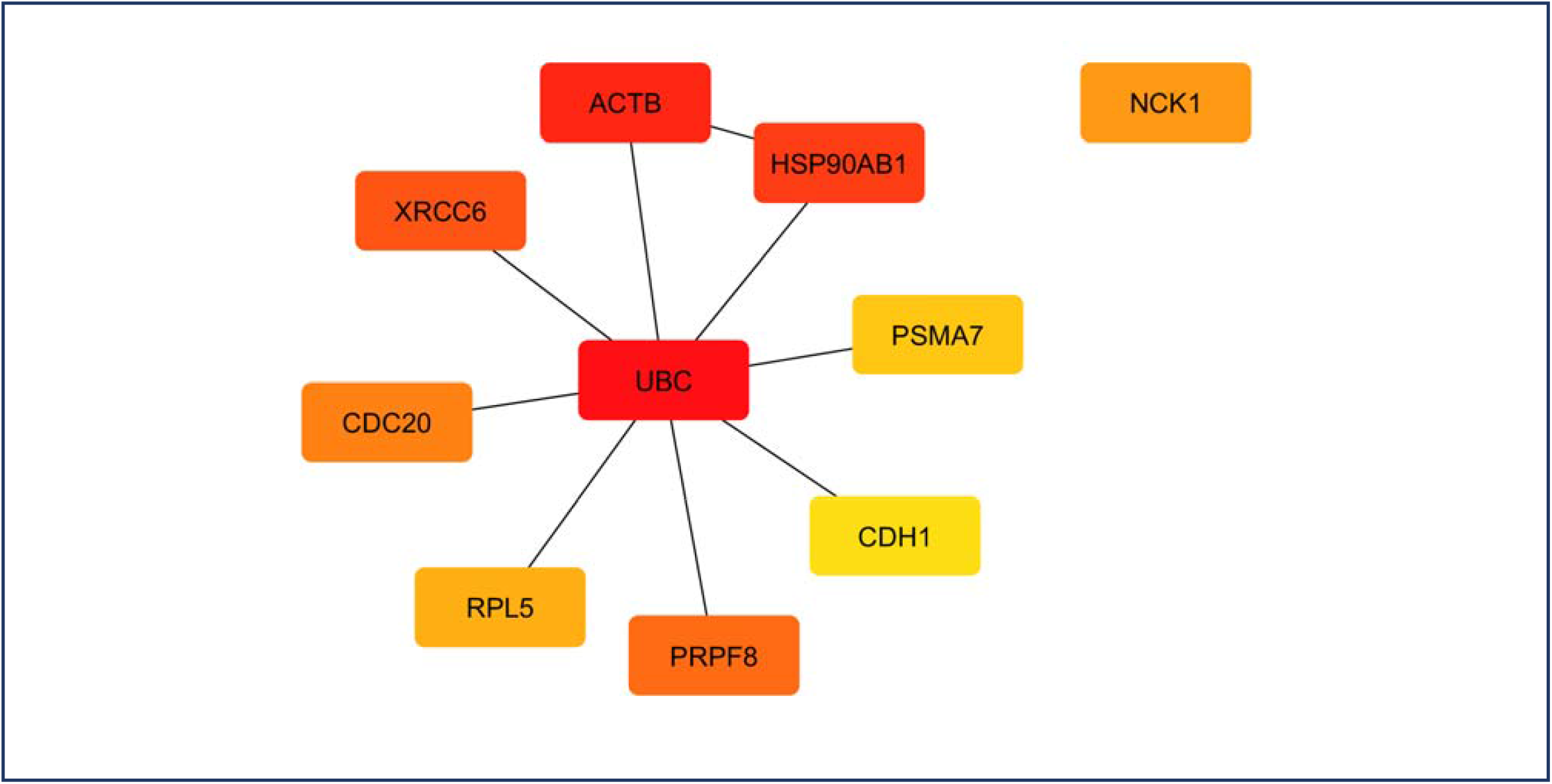
Selected hub proteins from the constructed generic PPI network. Nodes represent proteins and edges represent connection. The number of connections among nodes reduce in respect to reduction of color intensity.

**Table 3:** Summary of the identified hub proteins from PPI network.

| **Protein Name** | **Description** | **Functional Category** | **Clinical Significance** | **References** |
| --- | --- | --- | --- | --- |
| NCK1 | NCK Adaptor Protein 1 | Adaptor protein | Involved in Astrocytoma. | [56] |
| UBC | Ubiquitin C | Ubiquitylation protein | Involved in both AD and PD. | [57] [58] |
| ACTB | Beta-Actin | Cytoskeletal protein | Involved in AD. | [59] |
| PSMA7 | Proteasome 20S Subunit Alpha 7 | Proteasome subunit | Downregulated in PD. | [60] |
| CDH1 | Cadherin-1 | Cadherin protein | Downregulated in AD and other neurodegenerative diseases | [61] |
| PRPF8 | Pre-mRNA-processing-splicing factor 8 | mRNA splicing protein | Involved in retinitis pigmentosa and leukemia. | [62][63] |
| RPL5 | Ribosomal Protein L5 | Ribosomal subunit | Involved in the pathogenesis of multiple sclerosis. | [64] |
| CDC20 | Cell-division cycle protein 20 | Cell cycle regulatory protein | Differentially expressed in different forms of gliomas. | [65] [66] |
| XRCC6 | X-Ray Repair Cross Complementing 6 | DNA binding protein | Involved in the pathogenesis of autism spectrum disease. | [67] |
| HSP90AB1 | Heat Shock Protein 90 Alpha Family Class B Member 1 | Chaperone protein | Involved in different forms of cancers. | [68] |

**Table 4:** Different pathways associated with hub proteins selected from DEGs.

| **Pathway ID** | **Pathway** | **Adjusted P Value** | **Associated Genes** |
| --- | --- | --- | --- |
| P00004 | Alzheimer's disease-presenilin pathway | 0.001064 | CDH1, ACTB |
| P00012 | Cadherin signaling pathway | 0.002417 | CDH1, ACTB |
| P00044 | Nicotinic acetylcholine receptor signaling pathway | 0.03349 | ACTB |
| P00016 | Cytoskeletal regulation by Rho GTPase | 0.03446 | ACTB |
| P00053 | T cell activation | 0.03591 | NCK1 |
| P00049 | Parkinson's disease | 0.03978 | PSMA7 |

### 3.3. Regulatory Signatures in AD and PD

The identified hub genes were then analyzed for their connection with TFs and miRNAs in order to identify transcriptional (TFs) and posttranscriptional (miRNAs) regulatory signatures. YY1, NFKB1, BRCA1, TP53, GATA2, SREBF2, E2F1, FOXC1, RELA and NFIC were selected a the best interacting TFs with the hub genes from the TFs-hub gene interaction network (**Figure 4 and Table 5**). hsa-mir-186-5p, hsa-mir-92a-3p, hsa-mir-615-3p, hsa-let-7c-5p, hsa-mir-100-5p, hsa-mir-93-3p, hsa-mir-5681a, hsa-mir-484, hsa-mir-193b-3p and hsa-mir-16p-5p were selected as the best interacting microRNAs from miRNA-hub gene interaction map (**Figure 5 and Table 6**).

**Figure 4:**
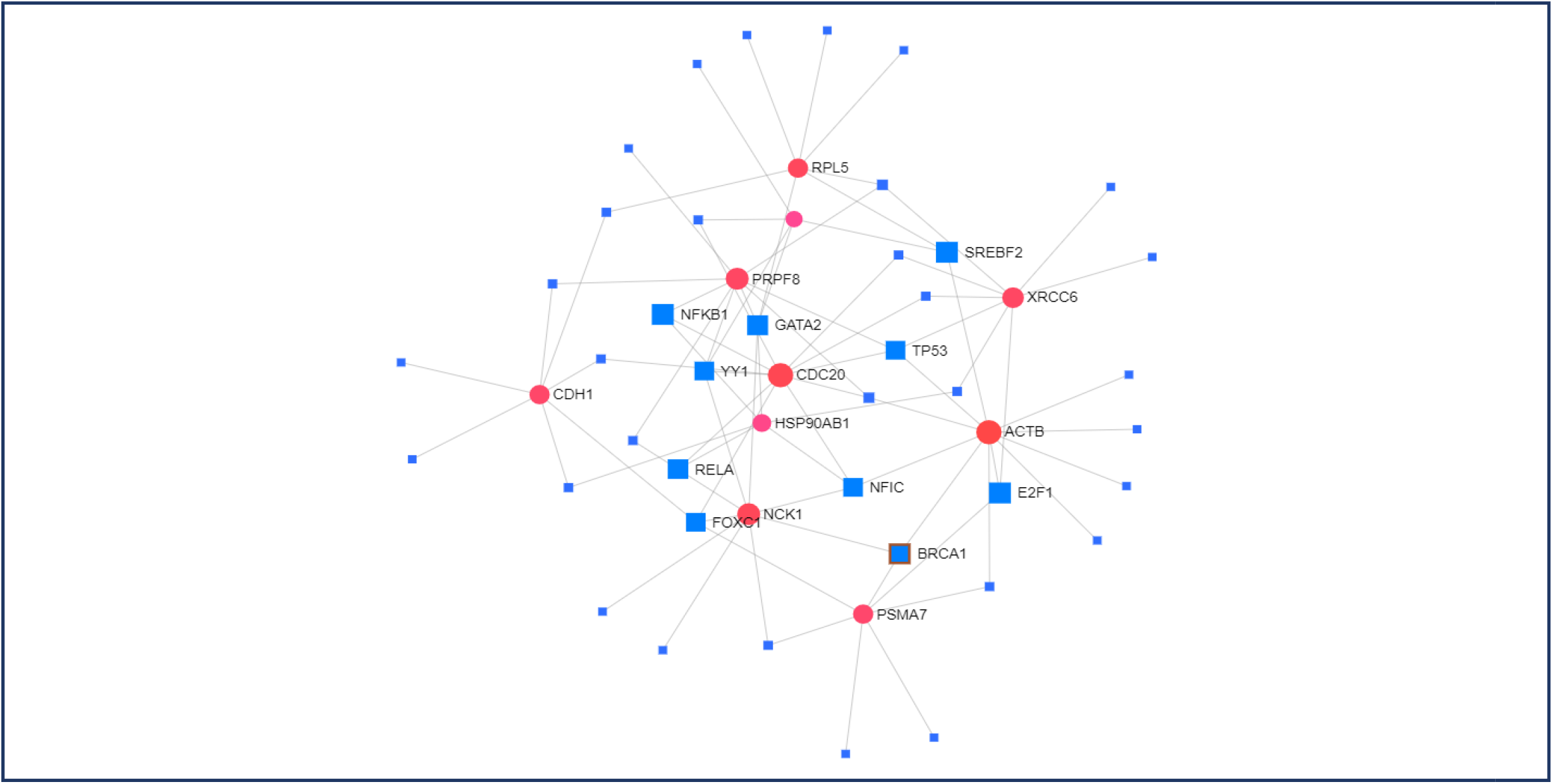
Hub gene-TFs interaction network. Nodes represent hub genes or TFs (Red: hub genes; Blue: Transcription factors) and edges represent interaction.

**Table 5:** Summary of the selected TFs from TFs-hub gene interaction network.

| TF Name | Description | Clinical Significance | References |
| --- | --- | --- | --- |
| YY1 | Yin Yang 1 | Downregulated in PD patients and leads to Aβ accumulation in AD | [69][70] |
| NFKB1 | Nuclear Factor NF-kappa-B p105 subunit | Highly expressed in AD patients. Mutated form was observed in PD | [71][72] |
| GATA2 | GATA binding factor 2 | Knockdown induces reduced expression of neuroglobin. Highly expressed in PD patients | [73][74] |
| SREBF2 | Sterol Regulatory Element Binding transcription factor 2 | Downregulated in AD and associated with bipolar disorder | [75][76] |
| E2F1 | Transcription factor E2F1 | Involved in AD.<br>Downregulated in PD | [77][78] |
| BRCA1 | Breast Cancer susceptibility protein | Reduced expression in AD | [79] |
| FOXC1 | Forkhead Box C1 | Reduced expression in PD patients and FOXC1 mutation is associated with age-related cognitive decline | [80][81] |
| TP53 | Tumour protein p53 | Associated with the pathogenesis of PD.<br>Involved in AD | [82][83] |
| RELA | Transcription factor p65/ Nuclear factor NF- kappa- B p65 subunit | Expression product induces inflammation in AD that contributes to the pathogenesis of early PD rats | [84][85] |
| NFIC | Nuclear factor I C | NFIC acts as a novel locus in AD | [86] |

**Figure 5:**
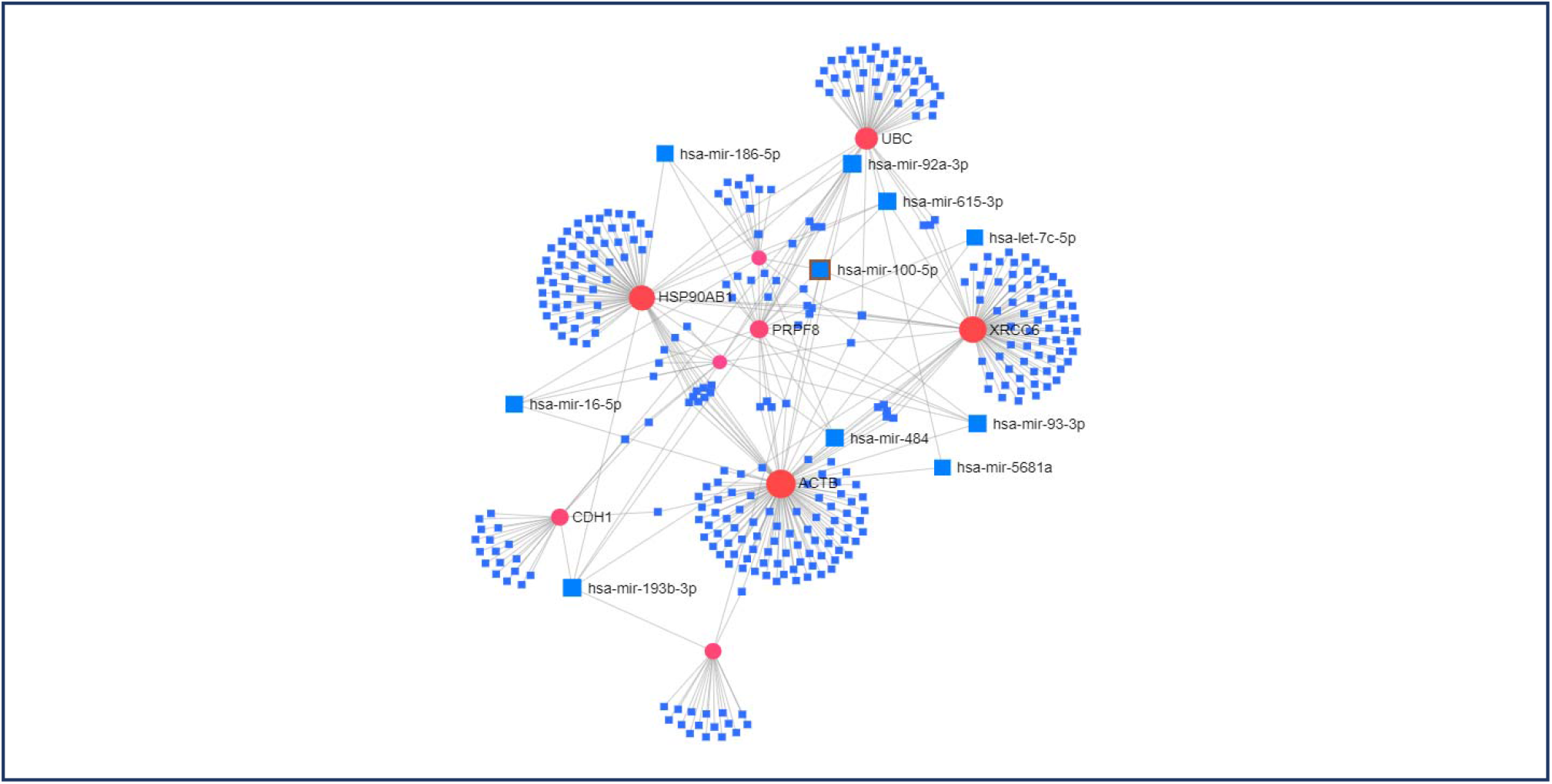
Hub gene-miRNAs interaction network. Nodes represent hub genes or miRNAs (Red: hub genes; Blue: miRNAs) and edges represent interaction.

**Table 6:** Summary of the selected miRNAs from miRNA-hub gene interaction network.

| miRNAs | Description | Clinical Significance | References |
| --- | --- | --- | --- |
| hsa-miR-186-5p | *Homo Sapiens* micro-ribonucleic acid -186-5p | Involved in Alzheimer's Disease (AD) and Multiple Sclerosis (MS) | [87][88] |
| hsa-miR-92a-3p | *Homo Sapiens* micro-ribonucleic acid -92a-3p | Involved in Alzheimer's Disease and Parkinson's Disease (PD) | [89]-[91] |
| hsa-miR-615-3p | *Homo Sapiens* micro-ribonucleic acid - 615-3p | Involved in Parkinson's Disease and Huntington's Disease (HD) | [92][93] |
| hsa-let-7c-5p | *Homo Sapiens* lethal -7c-5p | Involved in Parkinson's Disease | [94] |
| hsa-miR-100-5p | *Homo Sapiens* micro-ribonucleic acid -100-5p | Involved in Parkinson's Disease and Narcolepsy | [95] |
| hsa-miR-93-3p | *Homo Sapiens* micro-ribonucleic acid -93-3p | Involved in HIV-Associated Neurocognitive Disorder (HAND) | [96] |
| hsa-miR-5681a | *Homo Sapiens* micro-ribonucleic acid -5681a | Involved in Kidney Renal Clear Cell Carcinoma (KIRC) | [97] |
| hsa-miR-484 | *Homo Sapiens* micro-ribonucleic acid -484 | Involved in Alzheimer's Disease and Parkinson's Disease | [98][99] |
| hsa-miR-193b-3p | *Homo Sapiens* micro-ribonucleic acid -193b-3p | Involved in Alzheimer's Disease | [100] |
| hsa-miR-16-5p | *Homo Sapiens* micro-ribonucleic acid -16-5p | Involved in Alzheimer's Disease | [101] |

### 3.4. Epigenome Signatures in AD and PD

Hub genes were then analyzed for their histone modification sites to understand their mode of epigenomic modification. All of the selected hub genes were predicted to have both methylation and acetylation in different position except UBC which showed only sign of histone methylation (**Table 7**).

**Table 7:**
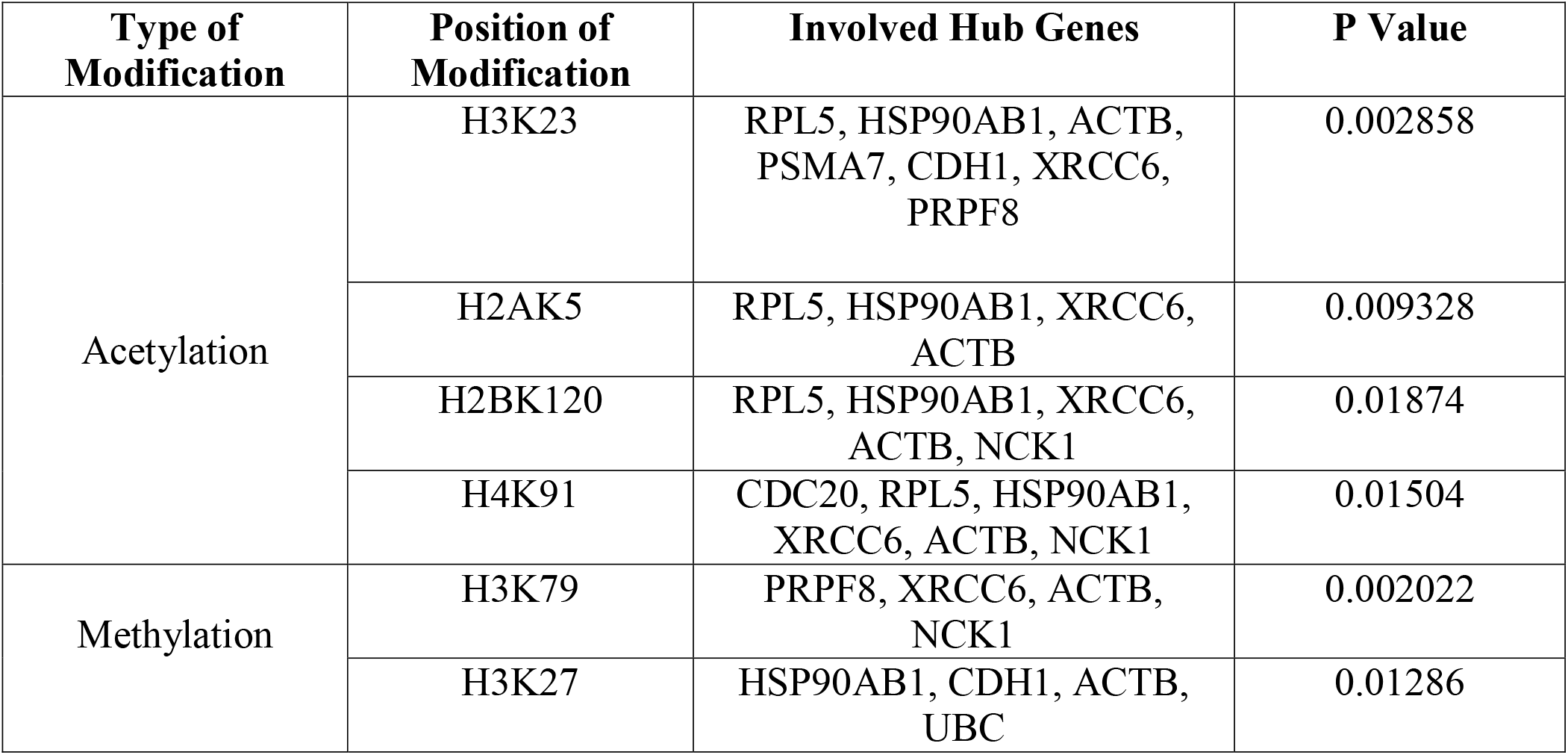
Pattern of histone modification in different hub genes involved in AD and PD.

| Type of Modification | Position of Modification | Involved Hub Genes | P Value |
| --- | --- | --- | --- |
| Acetylation | H3K23 | RPL5, HSP90AB1, ACTB, PSMA7, CDH1, XRCC6, PRPF8 | 0.002858 |
|  | H2AK5 | RPL5, HSP90AB1, XRCC6, ACTB | 0.009328 |
|  | H2BK120 | RPL5, HSP90AB1, XRCC6, ACTB, NCK1 | 0.01874 |
|  | H4K91 | CDC20, RPL5, HSP90AB1, XRCC6, ACTB, NCK1 | 0.01504 |
| Methylation | H3K79 | PRPF8, XRCC6, ACTB, NCK1 | 0.002022 |
|  | H3K27 | HSP90AB1, CDH1, ACTB, UBC | 0.01286 |

### 3.5. Identification of Therapeutic Candidate Molecules

All hub genes and genes encoding the TFs were analyzed to identify potential therapeutic candidate biomolecules based on gene-drug interaction. A number of drug molecules were identified which were predicted to act against a number of genes and thus may reverse AD and PD condition (**Table 8**). BRCA1 and TP53 were found to interact with most of the drugs with high drug scores. Antineoplastic agents were the most prominent drug category followed by kinase and proteasome inhibitors (**Figure 6A**). Among the identified drugs, 69% was found to be approved, 25% was in investigational stage and 6% was in the experimental phase (**Figure 6B**).

**Table 8:**
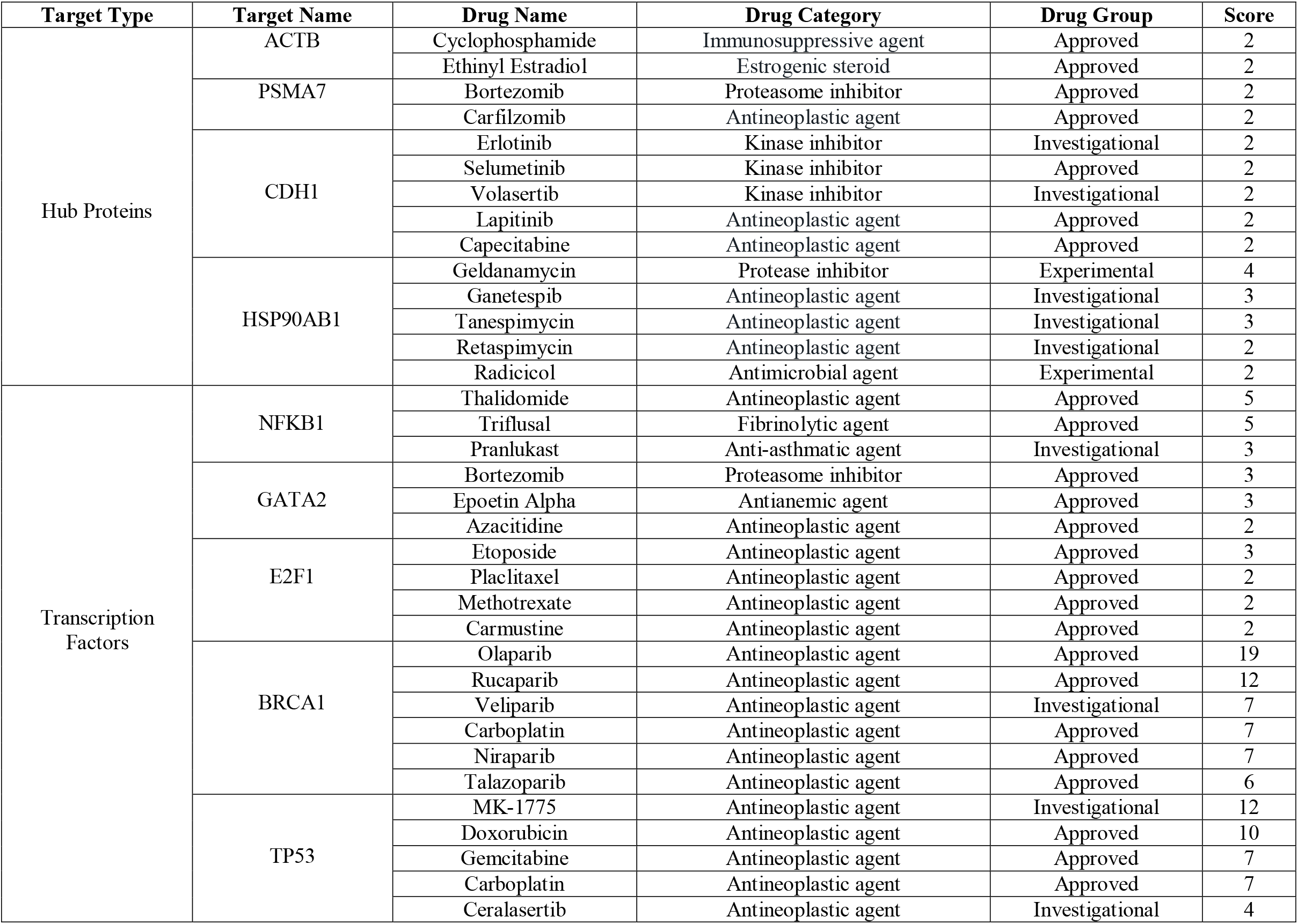
Identified candidate drug molecules based on drug-gene interaction expected to act against AD and PD.

**Figure 6:**
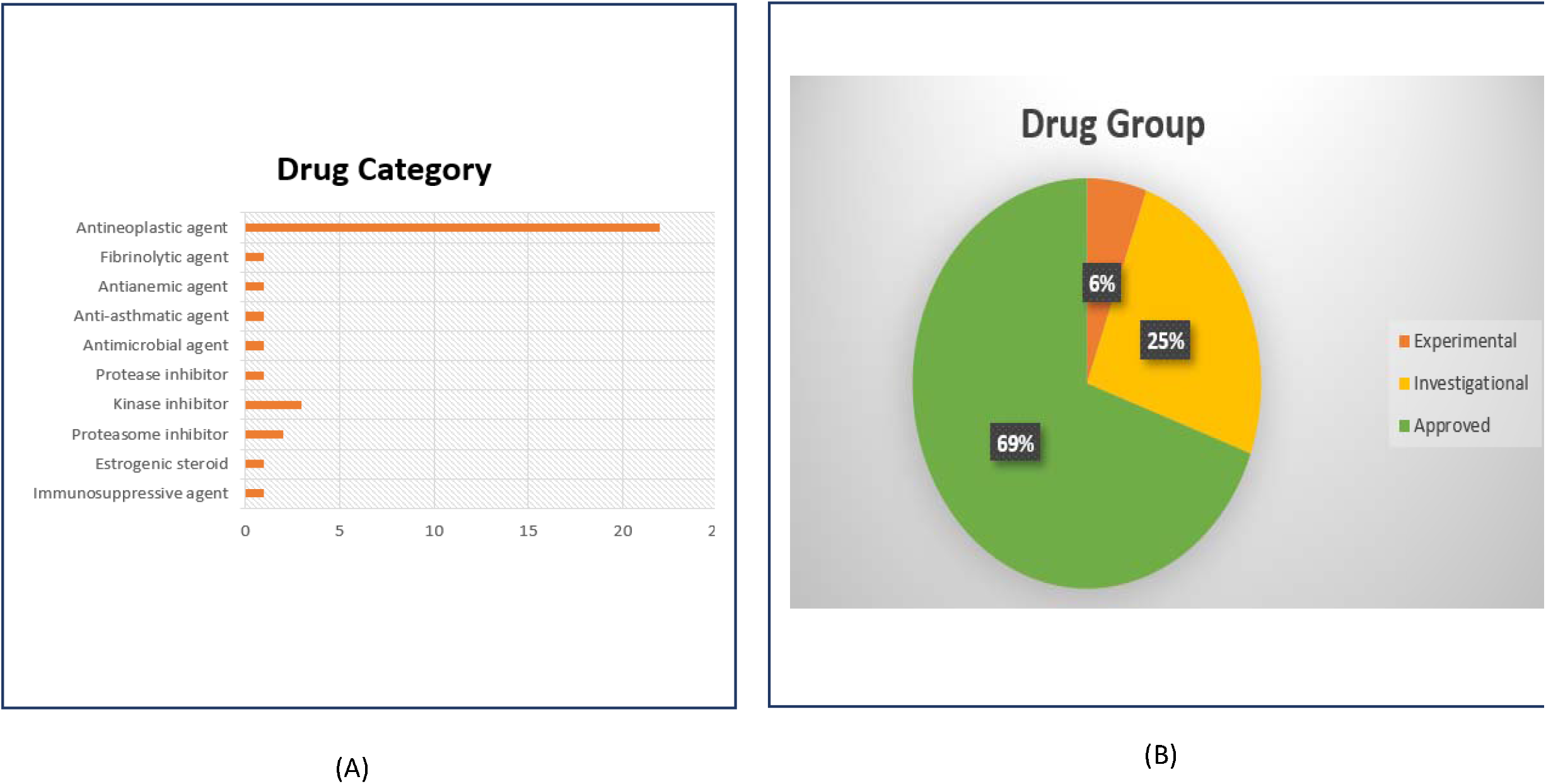
(A) Category; (B) Group classification of the identified drugs based on drug-gene interaction.

### 3.6. Subcellular Localization of the Hub Proteins

Only hub proteins were analyzed for their subcellular localization. Among the selected hub proteins, 50% was localized in cytoplasm, 40% in the nucleus and 10% was in plasma membran (**Figure 7**). UBC, PSMA7, ACTB, PRPF8 and HSP90AB1 were found to be localized in cytoplasm, CDC20, NCK1, RPL5 and XRCC6 were found operating in the nucleus and only CDH1 was reported to be located in plasma membrane.

**Figure 7:**
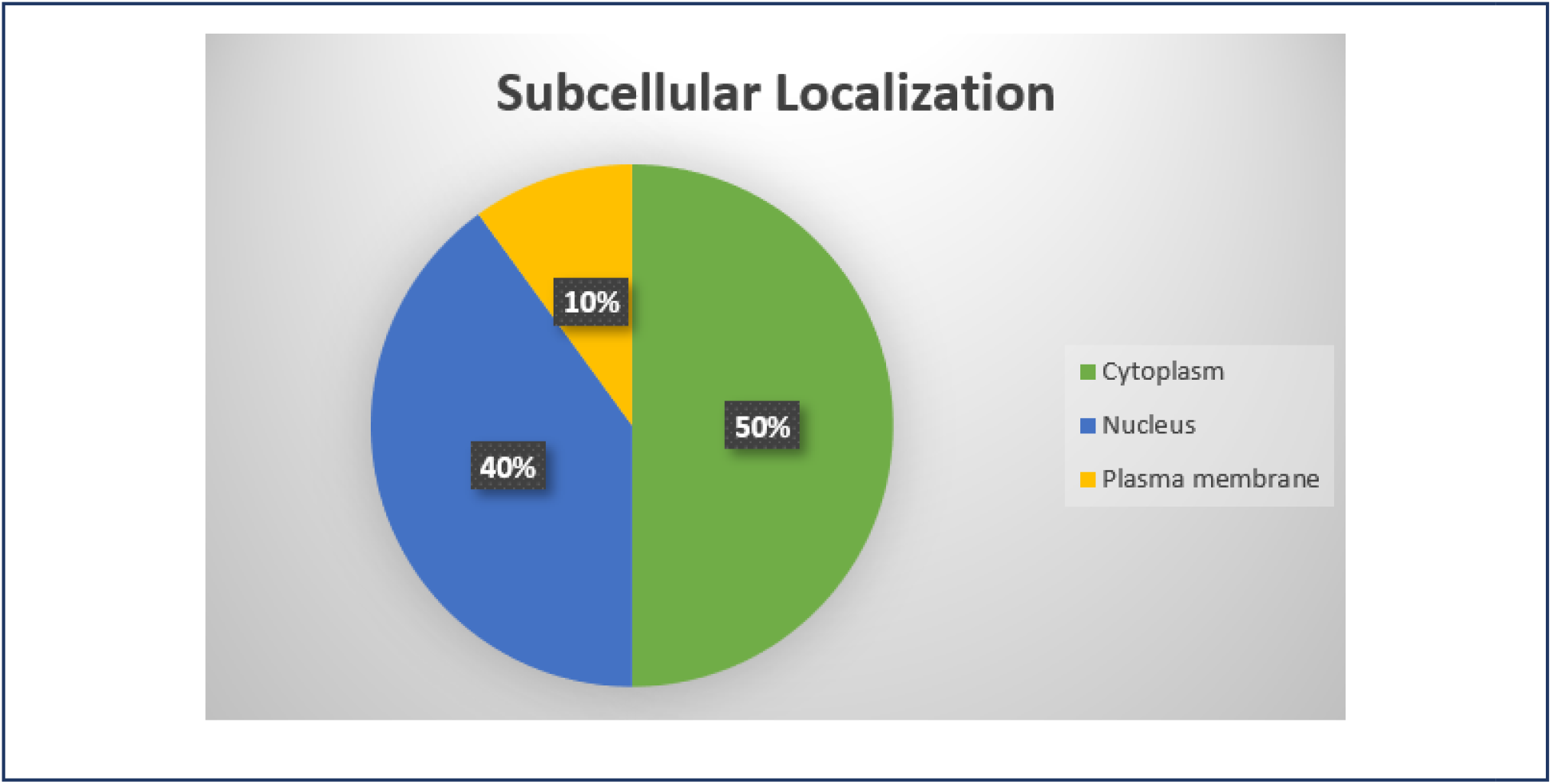
The distribution and percentile of subcellular localization shared by proteins expressed by the hub genes.

## 4. Discussion

In this study, publicly available microarray data of AD and PD patients were utilized in order to identify potential common biomarker signatures and molecular targets in both diseases. The two datasets were initially statistically analyzed to identify DEGs in both samples and then, the common DEGs of both datasets were taken for further experiments. Protein class overrepresentation analysis of the DEGs revealed that the DEGs were expressing mostly translational proteins (**Figure 1**). Gene ontology term and pathway analysis revealed that the DEGs were predominantly responsible for ribosomal function, protein processing, protein trafficking, ubiquitin-protein transferase inhibitor activity (**Table 1 and 2**). Protein translation is affected in both cases of AD and PD development as evident by the ribosome dysfunction in AD leading to decreased rate of protein synthesis and mRNA and tRNA level considered as the early event in AD prognosis. Moreover, different pattern of ribosomal protein phosphorylation is also encountered in case of PD [50][51]. Unusual protein processing i.e., abnormal cleavage of amyloid precursor protein (APP) leading to the formation of Aβ plaque, is a common phenomenon in AD. Missorting (protein trafficking) of microtubule-associated Tau protein is also associated in AD [52][53]. Dysregulated protein processing is also linked with the onset and progression of PD [54][55].

The DEGs were then analyzed to identify proteome signatures and mostly connected nodes (hub proteins) from the constructed generic PPI network (**Figure 3 and Table 3**). Pathway analysis of the hub proteins revealed the direct association of few hub genes with AD-presenilin pathway and PD (**Table 4**).

Among the selected hub proteins, a study revealed that, high level of expression of the NCK1 gene was involved in the prognosis of Astrocytoma, a form of malignancy of the brain. The study also suggested the NCK1 as the prognostic marker of Astrocytoma [56]. Along with Aβ and oxidative/nitrosative stress, a mutant form of ubiquitin gene called UBB^+1^, was found to be selectively expressed in the brains of AD patients while also impairing the proteasome activity *in vitro*. Induction/protection of the protein degradative system may be an efficient therapeutic strategy for neurodegenerative diseases such as AD and PD [57]. UBC is the central genome relating most genes in the PD networks and its role is related to protein synthesis, folding and degradation [58]. A laboratory experiment suggested that as ACTB had very low expression stability in the frontal cortex of Alzheimer’s patients thus is downregulated [59]. In a proteasome inhibition model of Parkinson’s disease, upon exposure to proteasome inhibitor lactacystin, microarray studies performed on primary cortical neurons revealed an upregulation of proteasome subunits such as PSMA7 [60]. Several findings have related to APC/C-Cdh1 downregulation to AD and showed that decrease in cdh1 protein levels are caused by both glutamate excitotoxicity and Aβ oligomers, leading to an accumulation of degradation targets and inactivation of the ubiquitin ligase. It has been shown that cyclin B1, PFKFB3, and glutaminase accumulate in AD due to APC/C-Cdh1 inactivation [61]. PRPF8 is essential for regulating mitophagy and its defects mediates the pathogenesis of neurodegenerative diseases. PRPF8 dysregulation is associated with the pathogenesis of retinitis pigmentosa [62]. Moreover, defective PRPF8 is responsible for miss-splicing of mRNA which eventually leads to leukemia [63]. Mutated form of RPL5 was found to be associated with the pathogenesis of multiple sclerosis, a severe disabling disease of the brain and spinal cord [64]. Appropriate centrosomal localization of CDC20 is crucial for the prompt development of the dendrites in the brain [65]. A laboratory experiment revealed both the upregulated and downregulated expression of CDC20 in gliomas and low-grade glioma tumors respectively [66]. A recent study has found different splicing patterns of XRCC6 in patients with autism spectrum disease (ASD) and suspected that reduced XRCC6 activity might be involved with the pathogenesis of ASD [67]. Downregulation of HSP90AB1 has been observed to be associated with breast and colorectal cancer (**Figure 3 and Table 3**) [68].

Then the hub-genes were analyzed for their interactions with transcription factors and miRNAs to identify transcriptional and post-transcriptional regulatory biomolecules. Among the selected TFs, YY1 provides neuroprotective function against SNCA toxicity. It is observed that in case of PD patients, YY1 is downregulated. Inhibition of YY1 signaling eventually leads to the degeneration of dopaminergic neurons, although the process of inhibition is yet to be determined [69]. YY1 in AD patients serves the function of an activator of the BACE1 promoter, one of the major β-secretase which assists in cleaving amyloid-beta peptide (Aβ) from APP. Accumulation of Aβ in the brain leads to the development of Alzheimer’s disease. YY1 can also maintain the level of Aβ by controlling the expression of molecules required for APP processing, e.g., FE65. Higher expression of FE65 corresponds to the development of severe AD [70]. Studies highlight higher expression of NFKB in AD patients. Immunoreactivity for p65 subunit was shown to be increased in neuronal elements specifically in the patient’s hippocampus and entorhinal cortex compared to the control group. AD is associated with the development of Aβ plaque which in turn regulates NFKB signaling. Activated NFKB induced higher expression of APP from which Aβ is cleaved [71]. A mutation analysis revealed several base changes in NFKB1 coding sequences of sporadic PD patients [72]. High expression of GATA2 is observed in Dopamine Cells and Brain Regions of a PD patient [73]. The knockdown of GATA2 was found to be involved in reduced expression of neuroglobin as evident in a laboratory experiment [74]. SREBF2 genes which are involved in cholesterol metabolism are found to induce AD prognosis. It was found that the mRNA level of SREBF2 was lower in AD patients compared to healthy individuals and thus it can be responsible for inducing the disease [75]. SREBF2 polymorphism is also associated with bipolar disorder [76]. E2F1 was found throughout the cytoplasm and neurites of PC12 cells in response to Aβ and in the cytoplasm of cells in the AD brains [77]. Upregulation of E2F1 is observed in PD patients [78]. Laboratory results showed reduction in neuronal BRCA1 in AD patients [79]. Reduced expression of FOXC1 was observed in the midbrain dopaminergic (DA) neurons of PD [80]. FOXC1 mutation influences cerebral small-vessel disease which is associated with age-related cognitive decline [81]. TP53 also known as p53 induces apoptosis and it has been shown through laboratory experiments the amount of p53 in the temporal cortex was significantly higher in AD brains than in the controls, and that immunoreactivity similar to p53 was observed in glial cells. [82]. TP53 is associated in many cell death pathways and its activation is involved in the pathogenesis of PD [83]. The transcription factor p65 of the NF-KB complex is expressed by RELA. High levels of NF-KB is found to be responsible for inducing inflammation in AD brains [84]. NF-Kb signaling was also recorded in the pathogenesis of early PD rats [85]. NFIC in Alzheimer’s patients was found as a novel locus (**Figure 4 and Table 5**) [86].

Among the selected miRNAs, hsa-miR-186-5p was found to be upregulated in the blood samples of patients with AD compared to normal controls, when reanalyzed from a pre-existing small RNA-Sequencing dataset [87] (**Table 6 and Figure 5**). Similarly, it was also found to be expressed in much higher levels in patients with multiple sclerosis, further confirming its association with other neurological disorders [88]. In a study, hsa-miR-92a-3p was found to be down-regulated in serums of PD patients and up-regulated in the cerebrospinal fluid (CSF) of AD patients [89] - [91]. Another study in comparison to previous studies revealed hsa-miR-615-3p to be a key regulator of PD as well as the up-regulator, in the prefrontal cortex of Huntington’s disease (HD) patients [92] [93]. A differential expression analysis has also revealed an association of hsa-let-7c-5p and hsa-miR-100-5p with PD [94]. The analysis of samples obtained from miRNA microarray in a study, with the help of qRT PCR, confirmed the down-regulation of hsa-miR-100-5p in each of the peripheral blood samples [95]. Moreover, an examination of miRNA brain profiles also revealed an up-regulation of hsa-miR-93-3p in HIV-Associated Neurocognitive Disorder (HAND) patients, which is known to repress peroxisome biogenesis factors leading to neurocognitive dysfunctions [96]. Another study which analyzed the differentially expressed genes for Kidney Renal Clear Cell Carcinoma (KIRC) found hsa-miR-5681a to be one of the targeting miRNAs for the KIRC genes in the network [97]. In a laboratory experiment with AD patients, the human brain hsa-mir-484 was found to be deregulated in the temporal cortex [98]. Again, in another study an association of the same miRNA with PD specific genetic pathway was revealed [99]. A meta-analysis study also revealed hsa-miR-193b-3p to be dysregulated in patients with both mild cognitive impairment (MCI) and Alzheimer’s Disease Dementia (ADD) [100]. hsa-miR-16-5p was also found to be deregulated (down-regulated) in several studies, in brain tissues of patients with Late-onset Alzheimer’s Disease (LOAD) [101].

Different types of epigenome signatures i.e., histone acetylation and methylation were also observed for hub genes which indicate their tight and controlled regulation and these changes may influence the contribution of these genes in disease induction (**Table 7**). Different therapeutic candidate molecules i.e., antineoplastic, proteasome inhibitor, kinase inhibitor was also identified upon screening the identified hub genes and TF encoding genes against drug database which may reverse the AD and PD condition (**Table 8**). Different identified drugs are in different stages of clinical development and few of them are already approved for medical use (**Figure 6**). Understanding the subcellular localization of a molecular target molecule is crucial for drug targeting. For instance, a drug targeting CDC20 needs to migrate into the nucleus. The hub proteins were analyzed for their subcellular localization and they were found to be distributed in cytoplasm, nucleus and plasma membrane (**Figure 7**).

Finally, this study recommends NCK1, UBC, CDH1, CDC20, ACTB, PSMA7, PRPF8, RPL7, XRCC6 and HSP90AB1 as the best proteome signatures, YY1, NFKB1, BRCA1, TP53, GATA2, SREBF2, E2F1, FOXC1, RELA and NFIC as the best transcriptional regulatory signatures and hsa-mir-186-5p, hsa-mir-92a-3p, hsa-mir-615-3p, hsa-let-7c-5p, hsa-mir-100-5p, hsa-mir-93-3p, hsa-mir-5681a, hsa-mir-484, hsa-mir-193b-3p and hsa-mir-16p-5p as the best post-transcriptional regulatory signatures based on the strategies employed in the overall study. The identified signatures could be used to identify potential biomarkers and drug targets for efficient diagnosis and treatments of AD and PD. The proposed networks, pathways and other analyses could also provide useful insight in the discovery of diagnostic and therapeutic interventions. And the identified candidate molecules could be investigated for possible drug discovery aiming to be triggered against both AD and PD.

However, the integrated network-based approach comes with few limitations i.e., mRNA level of a cell doesn’t always correlate with proteome level and this strategy can’t consider the post-translational modification. Apart from this, biomarkers for a particular disease is considered more feasible if it is secreted in bloodstream (secretome) [102]. So, these shortcomings need to be addressed by further *in vivo* and *in vitro* study which are currently underway.

## 5. Conclusion

AD and PD are two most prevalent forms of neurodegenerative disease which affect mass population around the world. Although, this disease is severely affecting a large number of world population but neither an efficient diagnosis nor an effective treatment options are currently available. In this study, two distinct microarray datasets for AD and PD were used in conjunction with network-based strategy to identify potential common biomarker signatures, drug targets and therapeutic agents for both AD and PD. Pathway and ontology analyses of the data revealed many significant correlations among different processes and disease progression. Network analysis of the key proteins suggested different proteome, transcriptional and post-transcriptional signatures which can be used in the discovery of common biomarkers and drug targets in AD and PD. Different candidate molecules were also identified which are expected to be used to reverse the AD and PD condition. However, since this experiment is largely based on computational study so, authors suggest further in vivo and in vitro study to strengthen the findings of this study.

## Declarations

### Ethics Approval and Consent To Participate

Not Applicable

### Consent for Publication

Not Applicable

### Availability of Data and Material

All the data are provided within the manuscript.

## Authors’ Contributions

NF conceived the study. NF, DP, SM and TR designed the study. NF, DP and SM wrote the draft manuscript. NF, AD, MA and TR edited and revised the manuscript. All the authors approved the final version of the manuscript.

## Acknowledgement

Authors are thankful to the members of Community of Biotechnology and Swift Integrity Computational Lab, Dhaka, Bangladesh for the supports during the preparation of the manuscript.

